# The miR156-targeted *SlSBP15* represses tomato shoot branching via modulating auxin transport and interacting with *GOBLET* and *BRANCHED1b*

**DOI:** 10.1101/2022.12.21.521468

**Authors:** Carlos Hernán Barrera-Rojas, Mateus Henrique Vicente, Diego Armando Pinheiro Brito, Eder M. Silva, Aitor Munoz Lopez, Leticia F. Ferigolo, Rafael Monteiro do Carmo, Carolina M. S. Silva, Geraldo F.F. Silva, Joao P. O. Correa, Marcela M. Notini, Luciano Freschi, Pilar Cubas, Fabio T.S. Nogueira

## Abstract

The microRNA156 (miR156)/*SQUAMOSA PROMOTER-BINDING PROTEIN-LIKE* (*SPL/SBP*) regulatory hub is highly conserved among phylogenetically distinct species, but how it interconnects multiple pathways to converge to common integrators controlling shoot architecture is still unclear. Here, we demonstrated that the miR156/*SlSBP15* hub modulates tomato shoot branching (SB) by connecting phytohormones with important genetic pathways regulating both axillary bud (AB) development and outgrowth. We verified that plants overexpressing the miR156 (156-OE plants) display high SB, whereas plants overexpressing a miR156-resistant SlSBP15 alelle (rSBP15 plants) display arrested SB and are able to partially restore the wild-type (WT) phenotype in156-OE background. Although rSBP15 plants showed ABs smaller than MT, its activation is dependent on shoot apex-derived auxin transport inhibition. Additionally, hormonal measurements reveal that IAA and ABA concentrations were lower in 156-OE and higher in rSBP15-OE plants. SlSBP15 regulates AB development and outgrowth by inhibiting auxin transport and the activity of *GOBLET* (*GOB*), and by interacting with BRANCHED1b (SlBRC1b) at the protein level to control abscisic acid (ABA) levels within ABs. Our data provide a new mechanism by which the miR156/*SPL/SBP* hub regulates SB, and suggest that *SlSBP15* has potential applications in improving tomato architecture.

## Introduction

Plant architecture is defined as the three-dimensional organization of the plant body (Reinhardt and Kuhlemeier, 2002). While the root apical meristem will form the root system, the shoot apical meristem (SAM) gives rise to the shoot architecture. The shoot architecture is determined by the organization and activities of apical, inflorescence, and axillary meristems (AM; Domagalska and Leyser, 2011). Once AM is established, it possesses the same developmental potential of the SAM by elongating and initiating few leaves in the axillary buds (ABs). ABs may either remain dormant or outgrow to develop lateral branches (LB), depending on a complex network involving both environmental and internal cues (Wang and Li, 2008; Domagalska and Leyser, 2011).

Several genetic pathways interact with phytohormones to control both AM establishment and AB outgrowth. Among the phytohormones, auxin, strigolactones, abscisic acid (ABA), and gibberellin (GA) act mostly as repressors, while cytokinin promotes shoot branching (SB; Dun *et al*., 2012). High levels of exogenous GA or ectopic GA biosynthesis in leaf axils reduces AM formation, impairing SB. Consistent with the fact that high levels of GA lead to the degradation of the GA-inhibitor DELLA, *Arabidopsis* quintuple *della* and tomato *della* (*procera*) mutants show impaired AM formation and inhibition of AB outgrowth (Hauvermale *et al*., 2012; Martínez-Bello *et al*., 2015; Zhang *et al*., 2020). ABA restricts AB outgrowth in *Arabidopsis* and promotes correlative growth inhibition under both high and low ratio of red light (R) to far-red light (FR) conditions (Reddy *et al*., 2013; Yao and Finlayson, 2015).

In order to outgrow ABs need to export auxin, which is limited by the strength of the AB, as an auxin source, and the sink strength of the polar auxin transport (PAT) stream in the main stem (Li and Bangerth, 1999; Balla *et al*., 2011). PAT stream is a mechanism mediated by the auxin efflux proteins PIN-FORMED (PIN) and ABCB (Gälweiler *et al*., 1998; Noh *et al*., 2001). Inhibiting PAT stream either by decapitating or treating the stem with auxin transport inhibitors triggers AB outgrowth (Domagalska and Leyser, 2011). While auxin, ABA, GA, and strigolactones inhibit AB outgrowth, CKs promote AB elongation in a dose-dependent manner, either when supplied directly to the ABs or stem (Wickson and Thimann, 1958; Dun *et al*., 2012; Roman *et al*., 2016), or overexpressing of CK biosynthesis-associated genes as observed in tobacco and tomato (Smigocki, 1991; Eviatar-Ribak *et al*., 2013, Pino *et al*., 2022).

In addition to phytohormones, the establishment and growth of AB depend on the action of several transcription factors. In tomato, for instance, AB requires *LATERAL SUPPRESSOR* (*LS*), a member of the VHIID protein family that modulates AM establishment (Rossmann *et al*., 2015; Schumacher *et al*., 1999). *LS* transcript accumulation in tomato depends on *GOBLET* (*GOB*) activity, which is another transcription factor crucial for AM initiation (Berger *et al*., 2009; Busch *et al*., 2011; Rossmann *et al*., 2015). *GOB* encodes a microRNA164 (miR164)-targeted NAC-domain family member, whose transcripts are localized in narrow stripes at the leaf margins, flanking the distal side of future leaflet primordia, and at the boundaries between the SAM and leaf primordia (Berger *et al*., 2009). While *gob* mutants are defective in AM formation, tomato plants harboring a miR164-resistant *GOB* version develop multiple ectopic lateral shoots (Raman *et al*., 2008; Busch *et al*., 2011), suggesting that *GOB* may control tomato AM initiation at least in part by regulating the expression of *LS* or by playing in similar genetic pathways (Rossmann *et al*., 2015).

Another fundamental modulator of the SB belongs to the TEOSINTE BRANCHED1/CYCLOIDEA/PCF (TCP) family of transcription factors (Cubas *et al*., 1999). *TEOSINTE BRANCHED1 (TB1)* or *BRANCHED1* (*BRC1*) and their orthologs are negative regulators of AB outgrowth (Doebley *et al*., 1997; Takeda *et al*., 2003; Aguilar-Martinez *et al*., 2007; Braun *et al*., 2012; Martín-Trillo *et al*., 2011; Liu *et al*., 2017). *Arabidopsis* BRC1 directly induces the expression of the homeodomain leucine zipper genes (*HD-ZIP*) *HOMEOBOX PROTEIN 21* (*HB21*), *HB40* and *HB53*. These proteins, together with BRC1, induce the expression of *9-CIS-EPOXICAROTENOID DIOXIGENASE 3* (*NCED3*), a key enzyme in ABA biosynthesis. As ABA acts as a negative regulator of AB activity (Yao and Finlayson, 2015), *BRC1* represses SB, at least in part, by inducing ABA accumulation within ABs (González-Grandío *et al*., 2017). In tomato, *BRANCHED1b* (*SlBRC1b*) retained the ancestral role of *TB1/BRC1-*like genes in AB arrest (Martín-Trillo *et al*., 2011).

In some species, the expression levels of *TB1/BRC1-like* genes are regulated by the SQUAMOSA PROMOTER-BINDING PROTEIN-LIKE (SPL/SBP) proteins (Xie *et al*., 2020; Liu *et al*., 2017). The SPL/SBP is a plant-specific transcription factor family, in which most members are targeted by the highly conserved microRNA156 (miR156), establishing the miR156/*SBP/SPL* regulatory hub. This regulatory hub has been associated to several aspects of plant development, acting by itself or interacting with phytohormones, affecting different processes such as phase transition, flowering, leaf patterning, fruit and root development (Wang and Wang, 2015; Barrera-Rojas *et al*., 2021; Barrera-Rojas *et al*., 2020; Yu *et al*., 2015; Silva *et al*., 2014; Silva *et al*., 2019). Low levels of *SPL/SBP* transcripts by overexpressing the miR156 are associated to higher SB in several species (Gou et al., 2017; Liu *et al*., 2017; Luo *et al*., 2012; Chuck *et al*., 2007; Schwab *et al*., 2005; Silva *et al*., 2014). In rice, the *IDEAL PLANT ARCHITECTURE1* (*IPA1, OsSPL14*) suppresses SB by directly activating the expression of *TB1/BRC1* (Lu *et al*., 2013; Liu *et al*., 2017; Xie *et al*., 2020). Other *SPLs/SBPs*, such as maize *UNBRANCHED3* and rice *OsSPL7* have been shown to suppress SB by modulating CK biosynthesis and auxin degradation, respectively (Du *et al*., 2017; Dai *et al*., 2018). Interestingly, low SB/tillering may have positive agronomic impacts on some crops, as exemplified by the significant increase in rice grain yield in field trials observed in the *IPA1* mutant (Jiao *et al*., 2010).

In tomato, low levels of the miR156-targeted *SlSBP13* transcripts triggers SB (Cui *et al*., 2020); however, the underlying mechanisms regarding this process are unknown. Here we show that the miR156-targeted *SlSBP15* represses tomato SB by modulating phytohormones responses and SB-associated genes, including *GOB* and *SlBRC1b*. Loss of miR156-targeted *SlSBP* function through miR156 overexpression (156-OE plants) increases SB and this effect is counteracted in plants that ectopically express a miR156-resistant version of *SlSBP15* (rSBP15 plants), the tomato homolog of *IPA1*. Genes associated with auxin transport and ABA biosynthesis were mis-regulated in ABs from 156-OE and rSBP15 plants, which is consistent with their distinct auxin and ABA levels. Although *SlSBP15* negatively regulates AB formation by repressing the activity of *GOB*, it is not sufficient to inhibit AM initiation and AB development in rSBP15 plants. On the other hand, the r*SBP15* is sufficient to repress SB in the gain-of-function *GOB* mutant. Importantly, we showed that SlSBP15 interacts with SlBRC1b at the protein level, and regulates common targets both cooperatively and opposingly. Taken together, these results provide a novel mechanism by which the miR156/*SlSBP15* hub interconnects complex networks to modulate tomato shoot architecture.

## Materials and methods

### Plant material and growth conditions

Tomato (*Solanum lycopersicum* L.) cultivar Micro-Tom LA3911 (MT) was used in this work otherwise is indicated. Transgenic MT plants overexpressing miR156 (156-OE) and the mutant *Goblet-4d* (*Gob-4d*) were previously described (Silva *et al*., 2014; Berger et al., 2009). Seeds were sowing and plants were grown under standard greenhouse conditions as described (Silva *et al*., 2019).

### Plant crosses, branching phenotyping, and Scanning Electron Microscopy (SEM)

For crosses, flowers were emasculated and manually pollinated 2 days before anthesis to prevent self-pollination. *Gob-4d* mutation was introgressed into the MT cultivar as described by Carvalho et al. (2011). Branching phenotype was evaluated as described (Busch et al., 2011) in 30-days post germination (-dpg) plants. Leaf axils were examined one by one beginning with the oldest leaf axil and annotated according to the following parameters: arrested/dormant ABs and lateral branches. For each experiment, at least seven plants of each genotype and crossing were analyzed. For SEM analysis, samples were fixed, mounted, and analyzed as described (Bharathan et al., 2002).

### Phylogenetic analysis

The SBP domains from distinct species were aligned using ClustalW (http://www.ebi.ac.uk/Tools/clustalw). Phylogenetic tree was constructed using the maximum likelihood algorithm with 1000 bootstraps replicates in MEGA7 (Kumar *et al*., 2016) using default parameters.

### *In Situ* hybridization

All steps were performed as described by Javelle *et al*. (2011). Shoot apexes from MT and 156-OE seedlings at 7- and 10-dpg were collected and fixed in 4% (w/v) paraformaldehyde. After alcoholic dehydration, shoot apexes were infiltrated and embedded in Paraplast X-Tra (McCormick Scientific). For *SlSBP15* probe preparation, pGEM vector containing a 597-bp fragment (nucleotides 523 to 1119 of the coding sequence) was linearized using the *BspHI* restriction enzyme (NEB). *In vitro* transcription was performed using the DIG RNA Labeling Kit (SP6/T7, Roche). Sense *SlSBP15* probe was used as a negative control. Locked nucleic acid (LNA) probe with sequence complementary to miR156 (5′-GTGCTCACTCTCTTCTGTCA-3′) was synthesized by Exiqon, and digoxigenin-labeled using a DIG oligonucleotide 3′ end labeling kit (Roche Applied Science). 10 picomols of each probe were used for each slide. All hybridization and washing steps were performed at 55°C. Pictures were photographed using light Microscope Axio Imager.A2 (Carl Zeiss AG). Oligonucleotide sequences are listed in Supplementary Table S1.

### RNA extraction, cDNA synthesis and qRT-PCR analysis

Total RNA was isolated using Trizol reagent (ThermoFisher) and treated with DNase I (Invitrogen). 1 μg of DNase I-treated RNA was used to generate the cDNA according to Varkonyi-Gasic *et al*. (2007). qRT-PCR reactions were performed using Platinum SYBR Green qPCR SuperMix UDG (Invitrogen) and analyzed in a Step-OnePlus real-time PCR system (Applied Biosystems). For each genotype, at least three biological samples with three technical replicates were used in the qRT-PCR analyses. *ACTIN4* was selected as internal control by initially analyzing three housekeeping genes, *ACTIN4* (Solyc04g011500), *BETA-TUBULIN 1* (Solyc04g081490) and *ALPHA-TUBULIN 3* (Solyc04g077020) as described by Exposito-Rodriguez *et al*. (2008). Relative expression (RE) levels of each gene were calculated using the 2^− ΔΔCT^ method (Livak and Schmittgen, 2001). Oligonucleotide sequences are listed in Supplementary Table S1.

### CRISPR-Cas9 gene editing

Two guide RNAs (gRNAs) targeting the *SlSBP15* second exon were combined by the PCR overlapping to obtain a 50-bp delection on the SBP domain. Protospacers sequences were selected using the CRISPR/Direct and the CCTop - CRISPR/Cas9 target online predictor (Naito et al., 2015; Stemmer *et al*., 2015). Protospacers were joined with the scaffold gRNA and the P2A/Csy4 splicing system under the *CmYLCV* promoter, using the pDIRECT_22C vector (Addgene plasmid # 91135) as a template. PCR products were combined into a single fragment, and cloned into pDIRECT_22C by Golden Gate reaction using the *SapI* and T4 ligase. *Agrobacterium tumefaciens* strain GV3101 harboring the pDIRECT_22C/*SlSBP15* gRNA recombinant vector was used to transform tomato cv. MT as described (Pino *et al*., 2010). T0 plants were initially PCR-genotyped for *Cas9* and, then, PCRs for genotyping and sequencing the *SlSBP15* deletion were made using primers flanking the protospacers. Sequences were confirmed by Sanger sequencing. Oligonucleotide sequences are listed in Supplementary Table S1.

### Vector constructs and plant transformation

*p35S::rSBP15* construct was generated by introducing PCR-mediated synonymous mutations into the miR156 binding site to prevent the miR156-directed *SlSBP15* cleavage; cDNA from MT leaves was used as a template, and PCR product was cloned in pENTR D-TOPO and subsequently recombined into pK7WG2.0 (Gateway System) using LR Clonase (ThermoFisher Scientific). Constructs for transactivation assays were generated as following: five fragments of the *GOB* promoter were individually cloned into pUPD2 vector and then ligated upstream to the firefly *LUCIFERASE* (*LUC*) and NOS terminator in the pDGBα2 vector (Sarrion-Perdigones *et al*., 2013). *SlPIN9* and *SlNCED1* promoters (2166 bp and 1568 bp, respectively) were cloned in front of the firefly *LUC* and CaMV35S terminator in the pICH86966 vector. *p35S::GFP-NLS* and *p35S::SlBRC1b* constructs were gifts from Dr. Michael Nicolas (CNB/CSIC, Spain). Full length cDNAs of *SlSBP15* and *SlBRC1b* were amplified by PCR and cloned into pDONR207. Subsequently, the constructs were introduced into the pAB117 (GFP fusion), pAB118 (mcherry fusion) and pAB119 (GFP-mcherry fusions) binary vectors. All oligonucleotide sequences used in the constructs are listed in Supplementary Table S1.

### Luciferase transactivation assays

Reporter and effector constructs were agroinfiltrated into 5-week-old *Nicotiana benthamiana* leaves. For *pGOB::LUC*, LUC activity was evaluated two days after agroinfiltration in leaves sprayed with D-Luciferin (Promega) using the NEWTON 7.0 CCD imaging system (Vilber). Relative LUC activity in leaves was quantified using the NEWTON 7.0 imaging software (https://www.vilber.com/newton-7-0/) with default parameters. At least 5 leaves from different *N. benthamiana* plants were used. For evaluating LUC activity from *pPIN9::LUC* and *pNCED1::LUC* constructs, leaf disks were collected 2-days after agroinfiltration and incubated with D-Luciferin. Luminescence was measured in a luminometer every 30 minutes, during at least 18 hours. Each leaf was treated as a biological sample, from which five leaf disks were collected as technical replicates.

### Yeast-two-hybrid assay

For yeast-two-hybrid assay, *SlSBP15, SlBRC1a* and *SlBRC1b* ORFs were cloned into pDONR207 through a BP clonase reaction, and recombined into respective vectors through a LR clonase reaction. *SlSBP15* ORF was cloned into pGBKT7-BD vector while *SlBRC1a* and *SlBRC1b* ORFs were cloned into pGADT7-AD. Yeast strain AH109 was grown overnight in YPDA medium, and competent cells were prepared and Binding domain and Activation domain vectors were co-transformed using the lithium acetate/single-stranded carrier DNA/polyethylene glycol method (Gietz and Woods, 2002). Transformants were selected in media lacking leucine and tryptophan (-LH). Strains were assayed for interaction in media lacking leucine, tryptophan and histidine (-LWH) in the presence or absence of 5mM 3-AT. Each row shows ten-fold serial dilutions of the indicated strain with three colonies.

### Hormonal measurements and NPA treatment

Endogenous indoleacetic acid (IAA) and abscisic acid (ABA) levels were determined by gas chromatography-mass spectrometry (GC-MS) in selected ion monitoring (SIM). Frozen samples (∼50 mg FW) were extracted and methylated as described in Cruz *et al*. (2018). Approximately 0.5 µg of the labeled ABA standard ([^2^H_6_]-ABA, OlChemIm Ltd.) and 0.25 µg of the labeled IAA standard ([^13^C_6_]-IAA, Cambridge Isotopes, Inc.) were added to each sample as internal standards. Analysis was performed on a gas chromatograph as described in Cruz *et al*. (2018). Ions with a mass ratio/charge (m/z) of 130 and 189 (corresponding to endogenous IAA); 136 and 195 (corresponding to [^13^C_6_]-IAA); 134, 162 and 190 (corresponding to endogenous ABA); and 138, 166 and 194 (corresponding to [^2^H_6_]-ABA) were monitored. For 1-Naphthylphthalamic acid (NPA) treatment, apexes were treated with 10 µL of 50µM NPA or Mock at 10-, 12-, 14- and 16-dpg (Pattison and Catalá, 2012). SB phenotype was evaluated at 30-dpg. NPA stock solution (100 mM) was previously prepared in dimethyl sulfoxide (DMSO). An equivalent concentration of DMSO was used to prepare the mock solution (0.05% v/v).

### Differentially expressed genes (DEGs) analysis

Early ABs were collected from MT, 156-OE and rSBP15 plants at 25-dpg, and immediately frozen in liquid nitrogen. Two total RNA replicates from each genotype were sent to construct RNA sequencing libraries (RNA-seq) and high throughput sequencing (Illumina NovaSeq platform) at Fasteris Co. Ltd (https://www.fasteris.com/en-us; Switzerland). Raw sequencing reads were cleaned by removing adaptor sequences and low-quality reads. The resulting high-quality reads were mapped to the tomato reference genome (Tomato Genome Consortium, 2012) (ITAG4.0) and transcripts were assembled, mapped and quantified using the Hisat2, StringTie and Ballgown packages with default parameters (Pertea et al. 2016). Read count normalization and differential expression analysis were normalized by FPKM (Fragments per Kilobase Million; Mortazavi et al., 2008) and statistical comparative analysis between MT and 156OE plants and between MT and rSBP15-overexpressing plants were performed in the Ballgown package of R (http://www.bioconductor.org/). DEGs in ABs from 156-OE and rSBP15 plants compared to MT, with adjusted P ≤ 0.05 and ratio ≥ 1.5, were filtered for further analysis.

DEGs were annotated based on SOLGENOMICS database (version SL4.0 and Annotation ITAG4.0), and *Arabidopsis thaliana* best protein blast hit from the TAIR database (Supplementary Tables S2-S4). Enriched GO terms for the DEG list were identified using the PANTHER Overrepresentation Test in the Gene Ontology Consortium database (Gene Ontology Consortium, 2021). SOLGENOMICS database was used as reference for the GO analysis. Fisher’s exact test with FDR correction with a cutoff of 0.05 was applied to determine enriched terms (Benjamini and Hochberg, 1995). Venn diagram was performed online at http://bioinfomatics.psb.ugent.be/webtools/Venn. Moreover, common DEGs between 156OE and rSBP15 plants with contrasting expression (log2 fold-change values) were selected and a heatmap was generated using the pheatmap package for R (Kolde, 2019). Raw sequence data from this study have been deposited in Gene Expression Omnibus (GEO) of NCBI under the accession number GSE212774.

### APB–FRET assays

FRET-AB assays Protein fusion binary vectors were agroinfiltrated in *N. benthamiana* leaves. 24-hours after infiltration, leaves were sprayed with 20 mM estradiol and 0.05% Silwet solution to induce gene expression. Assays were performed using the FRET-AP wizard of LAS-AF with the following parameters: 700 Hz acquisition speed; pinhole 111.5 mm; image format 512 × 512 pixels; 5X zoom. A 4 mm-diameter circular region of interest (ROI) was defined and photobleached with three repeated exposures (laser at 561 nm, 100% power level). FRET efficiency (E_FRET_) was calculated as the percentage of increase in donor fluorescence after acceptor photobleaching; E_FRET_= D_post_ - D_pre_/D_post_, where D_pre_ and D_post_ are the fluorescence intensity of the donor before and after photobleaching, respectively. D_pre_ and D_post_ were measured inside the 4mm defined ROI.

## Results

### Over-expressing the miR156 in tomato enhances SB and its target *SlSBP15* cluster with branching-associated genes

Several highly conserved genetic pathways are directly associated with SB, including the miR156-controlled pathway (Wang and Wang, 2015; Guo *et al*., 2008; Silva *et al*., 2014; Cui *et al*., 2020); however, how this regulatory pathway controls tomato vegetative architecture at genetic and molecular levels is unknown. In a previous work in our laboratory, we have found that tomato plants that over-express the miR156 in the Micro-Tom (MT) background (hereinafter referred as 156-OE plants) display enhanced SB index compared to wild-type (Silva *et al*., 2014) suggesting that the miR156-controlled genetic pathway modulates SB in tomato. Thus, we initially assessed the SB pattern in 156-OE plants by comparing the developmental stage of LB on the main shoot of plants at 30-days post germination (-dpg). 156-OE plants display not only nearly 100% of activated ABs but also, two-fold the number of LB compared to MT plants (Supplementary Fig. S1A, B). To examine whether the SB phenotype caused by over-expression of the miR156 is not cultivar-dependent, we outcrossed 156-OE plants with tomato cv. Moneymaker LA2706 (MM) and analysed the F1 offspring, in which the known recessive MT mutations are in heterozygosis (Martí *et al*., 2006). As we expected, overexpression of miR156 in the hybrid offspring led to a similar highly branched phenotype as observed in 156-OE plants in the MT background (Supplementary Fig. S1C). These data suggest that the miR156-regulated *SlSBP* genes negatively regulate the SB pattern in a cultivar-independent manner in tomato.

*SBP/SPL* genes have been described in several species as repressors of SB, including *SPL7/14/17* from rice (Dai *et al*., 2018; Jiao *et al*., 2010; Miura *et al*., 2010), *SPL3*/*17* from bread wheat (Liu *et al*., 2017), and *SPL9/10/15* from Arabidopsis (Schwarz *et al*., 2008; Gao *et al*., 2018). In tomato, there are 15 *SPL* genes (hereinafter called as *SlSBP* genes), 10 of which are post-transcriptionally regulated by miR156 (Salinas *et al*., 2012; Morea *et al*., 2016); among them, the *SlSBP13* was previously associated to SB (Cui *et al*., 2020), however, an in-depth study describing the underlying mechanisms by which the miR156-targeted *SlSBP* genes control tomato SB is lacking.

Therefore, to initially identify other possible tomato orthologs of SB-associated *SBP* genes, we have performed a phylogenetic analysis by comparing the SBP-box domains from *Arabidopsis* (AtSPL), rice (OsSPL), bread wheat (TaSPL), and tomato (SlSBP) proteins. SPL/SBP proteins were grouped into seven clusters, each one containing at least one SlSBP protein. Among them, the *SlSBP15* (Solyc10g078700) grouped into the cluster containing known SB-associated SPL genes (Supplementary Fig. S1D), which suggests that the *SlSBP15* could negatively regulate SB in tomato.

### The miR156-dependent *SlSBP15* regulation is crucial for the establishment of tomato SB

Our phylogenetic analysis suggests that the *SlSBP15* modulates SB; thus, to begin unraveling the role of the miR156/*SlSBP15* hub in controlling tomato SB, we initially decided to evaluate, by *in situ* hybridization, the spatiotemporal expression pattern of mature miR156 and *SlSBP15* transcripts in shoot apexes of MT and 156-OE seedlings at 7- and 10-dpg (Supplementary Fig. S2). As we expected, strong miR156 signal was detected in the developed AMs as well as in the SAM of MT seedlings at 7-dpg (Supplementary Fig. S2A, B). Interestingly, a strong *SlSBP15* signal was detected in the leaf axils of MT, in which no AM was visible (Supplementary Fig. S2C); in contrast, weak signals of *SlSBP15* transcripts were detected in AMs and inactive ABs from MT seedlings at 7- and 10-dpg, which negatively correlated with the presence of miR156 transcripts (Supplementary Fig. S2D). In addition, no *SlSBP15* transcripts were detected in active ABs from 156-OE plants, though a weak signal was observed in the procambium (Supplementary Fig. S2E). No signal was observed using the *SlSBP15* sense probe (Supplementary Fig. S2F). These results indicate that SlSBP15 transcripts are negatively regulated by miR156 during.

Our spatiotemporal expression data suggest that the *SlSBP15* inhibits SB in tomato, affecting the AM and AB development; thus, to confirm this hypothesis, we have generated both gain- and loss-of-function lineages of the *SlSBP15*. Gain-of-function lineage was obtained by introducing PCR-mediated synonymous mutations into the miR156 binding site to prevent the miR156-directed *SlSBP15* cleavage (Fig. 1A), here referred as miR156-cleavage resistant *SlSBP15* plants (rSBP15); loss-of-function lineage of the *SlSBP15* was obtained by employed the clustered regularly interspaced short palindromic repeats (CRISPR)-*ASSOCIATED NUCLEASE 9* (Cas9) gene-editing technology to generate null *slsbp15* mutant (here referred as *sbp15*^*CRISPR*^; Fig. 1B); then, we have evaluated the SB pattern in rSBP15 and *sbp15*^*CRISPR*^ lineages by comparing the developmental stage as previously described. Given all rSBP15 lines displayed similar *SlSBP15* expression levels (Supplementary Fig. S3A), we selected line #30 for further analyzes. rSBP15 plants were smaller than MT and almost all its ABs remained at the early/dormant developmental stage at 30-dpg (Fig. 1C, D; Supplementary Fig. S3B, C). *sbp15*^*CRISPR*^ plants were also smaller than MT, although they showed no obvious alterations in the SB pattern (Figure 1D-E).

**Figure 1.**
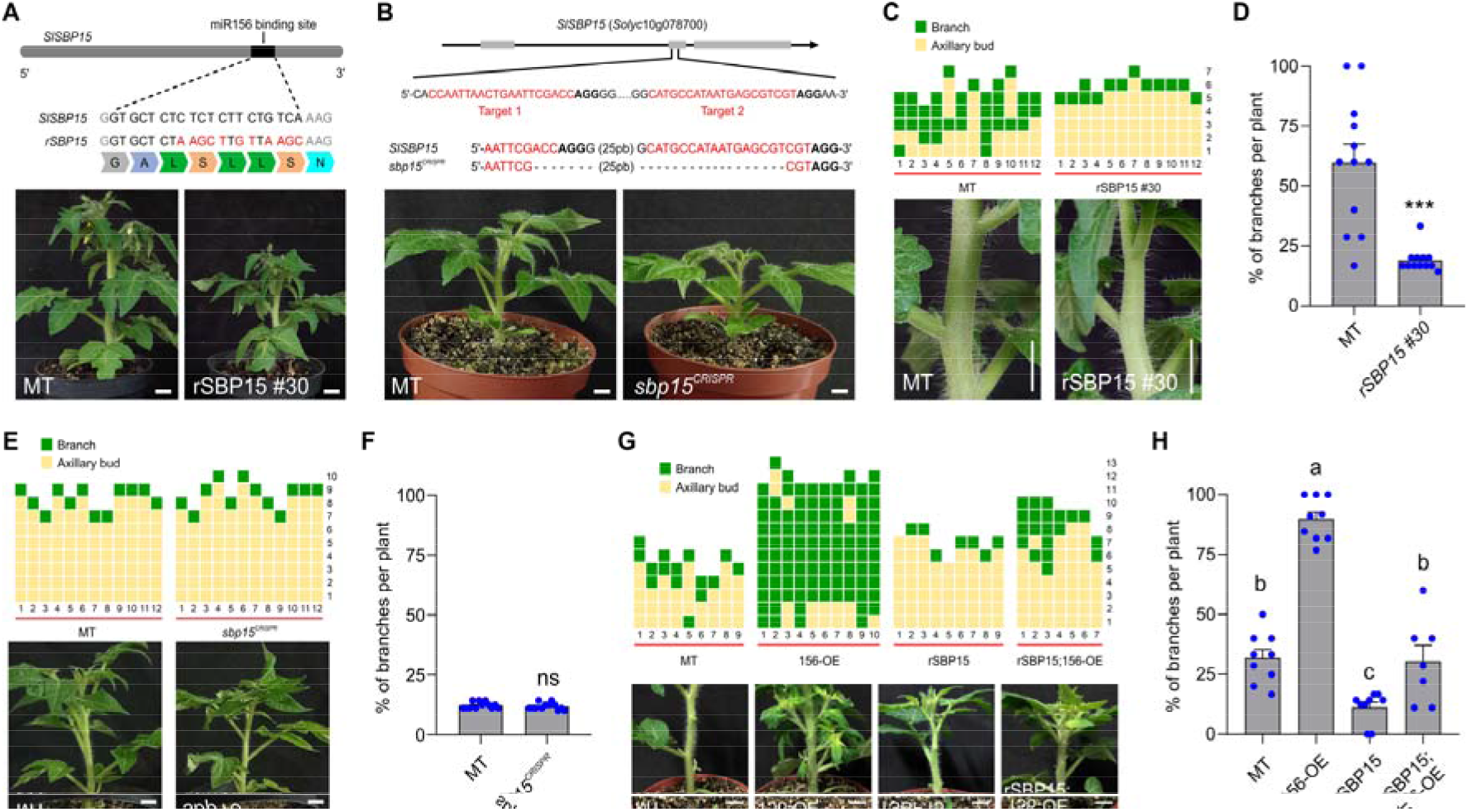
De-repression of *SlSBP15* suppresses tomato SB. (A) Schematic representation of *SlSBP15* CDS (top). Grey boxes represent CDS, black box represents miR156 binding site. Synonymous mutations modified at the miR156 binding site to prevent cleavage and produce a resistant *SlSBP15* allele (rSBP15). Representative pictures of MT and rSBP15 plants at 30-dpg (bottom). Scale bars = 1 cm. (B) (Top) The second exon of *SlSBP15* was targeted with two single-guide RNAs (sgRNA; target 1 and target 2 are highlighted in red font) for CRISPR/Cas9-induced gene editing. Bold fonts indicate the Protospacer-Adjacent Motif (PAM) sequences. (Middle) DNA sequencing analysis showed that *sbp15*^*CRISPR*^ allele harbors a deletion of 50 pb. The sequence gap length is shown in parentheses. (Bottom) Representative pictures of MT and *sbp15*^*CRISPR*^ plants at 30-dpg. Scale bars = 1 cm. (C) SB pattern (top) and representative pictures (bottom) of MT and rSBP15 plants at 30-dpg. Scale bars = 1 cm. Each column represents a single plant and each square indicates the status of each leaf axil starting from the first node above the cotyledons. ABs (orange squares) or LB (dark green squares). (D) Percentage of LBs per plant in MT and rSBP15 genotypes, at 30-dpg. The percentage was determined based on the number of leaf axils that present LBs. Values are mean ± SEM (n = 12). \*\*\**P*<0.001, according to Mann Whitney test. (E) *sbp15*^*CRISPR*^ showed a SB pattern similar to MT. SB pattern (top) and representative pictures (bottom) of MT and *sbp15*^*CRISPR*^ plants at 30-dpg. Scale bars = 1 cm. (F) Percentage of LBs per plant in MT and *sbp15*^*CRISPR*^ genotypes, at 30-dpg. Values are mean ± SEM (n = 12). ns: not significant. (G) *rSBP15* allele represses SB in the 156-OE background. SB patterns (top) and representative pictures (bottom) of MT, 156-OE, rSBP15, and rSBP15;156-OE plants at 30-dpg. Scale bars = 1 cm. (F) Percentage of LBs per plant in MT, 156-OE, rSBP15, and rSBP15;156-OE genotypes. Values are mean ± SEM (n = 7-10). Distinct letters represent statistical differences at 5% level evaluated by Tukey’s HSD test. The blue dots represent individual data points.

SB phenotypes from gain- and loss-of-function lineages indicate that the miR156-dependent regulation of *SlSBP15* is important for AB outgrowth, although *SlSBP15* can act redundantly with other *SlSBPs* to establish tomato vegetative architecture. To confirm this information, we have outcrossed the rSBP15 plants with 156-OE plants, and have analyzed the SB phenotype. The miR156-resistant version of the *SlSBP15* into the 156-OE background (rSBP15;156-OE plants) partially restored WT-like SB phenotype (Fig. 1G, H), indicating that the *SlSBP15* acts redundantly with other *SlSBP* genes in tomato SB, most likely *SlSBP13* (Cui *et al*., 2020). Furthermore, to examine whether the SB phenotype caused by the resistant version of the *SlSBP15* is also not cultivar-dependent, we have outcrossed rSBP15 plants with tomato MM background and have analysed the F1 offspring. As we expected, the resistant version of the *SlSBP15* into the hybrid offspring lead to inhibited AB outgrowth compared to the control F1 offspring, similarly to rSBP15 plants (Supplementary Fig. S3D). These data suggest that the de-regulation of the *SlSBP15* negatively regulate the SB pattern in a cultivar-independent manner in tomato.

### ABs from rSBP15 plants are functionally dependent on loss of PATs

Our data have shown that ABs from rSBP15 plants remained at the early/dormant developmental stage; also, they were smaller or less developed than those from MT (Fig. 2A), which suggests that rSBP15 plants could have not functional ABs. To answer this question, we decided to evaluate the ABs response from rSBP15 plants to decapitation, a classical assay to evaluate AB activation after release from apical dominance (Thimann and Skoog, 1933); thus, we have removed the SAM from rSBP15 plants at 30-dpg, and have measured the AB outgrowth in nodes 1, 2, and 3. All ABs from rSBP15 plants were able to outgrow in response to decapitation, but at distinct levels, compare to MT (Fig. 2B, C). Seven days after decapitation, the size of ABs at node 3 increased 135% in MT, relative to ABs in intact plants, whereas ABs in decapitated rSBP15 plants increased 800%. Fourteen days after decapitation, the most apical AB in MT and rSBP15 plants usually dominated over the others; however, in rSBP15 plants, the dominance from the topmost AB was stronger than in MT, as it grew nearly 3 times longer than the lower ABs (Fig. 2C). These observations indicate that rSBP15 plants have functional ABs and suggests that the loss of PAT triggers ABs activation. To confirm whether PAT is associated with the AB inhibition observed in rSBP15 plants, we have evaluated AB outgrowth in rSBP15 plants at 30-dpg treated with 1-Naphthylphthalamic acid (NPA), a well-known PAT inhibitor (Bennett *et al*., 2006). Interestingly, NPA-treated rSBP15 plants displayed active ABs between nodes 1 to 3, but the most upper AB in rSBP15 plants remained dormant compared to MT (Fig. 2D, E). These results are consistent with our decapitation experiments, reinforce that ABs from rSBP15 plants are functionally dependent on shoot apex-derived auxin transport inhibition, and also suggest that *SlSBP15* is involved in modulating auxin responses.

**Figure 2.**
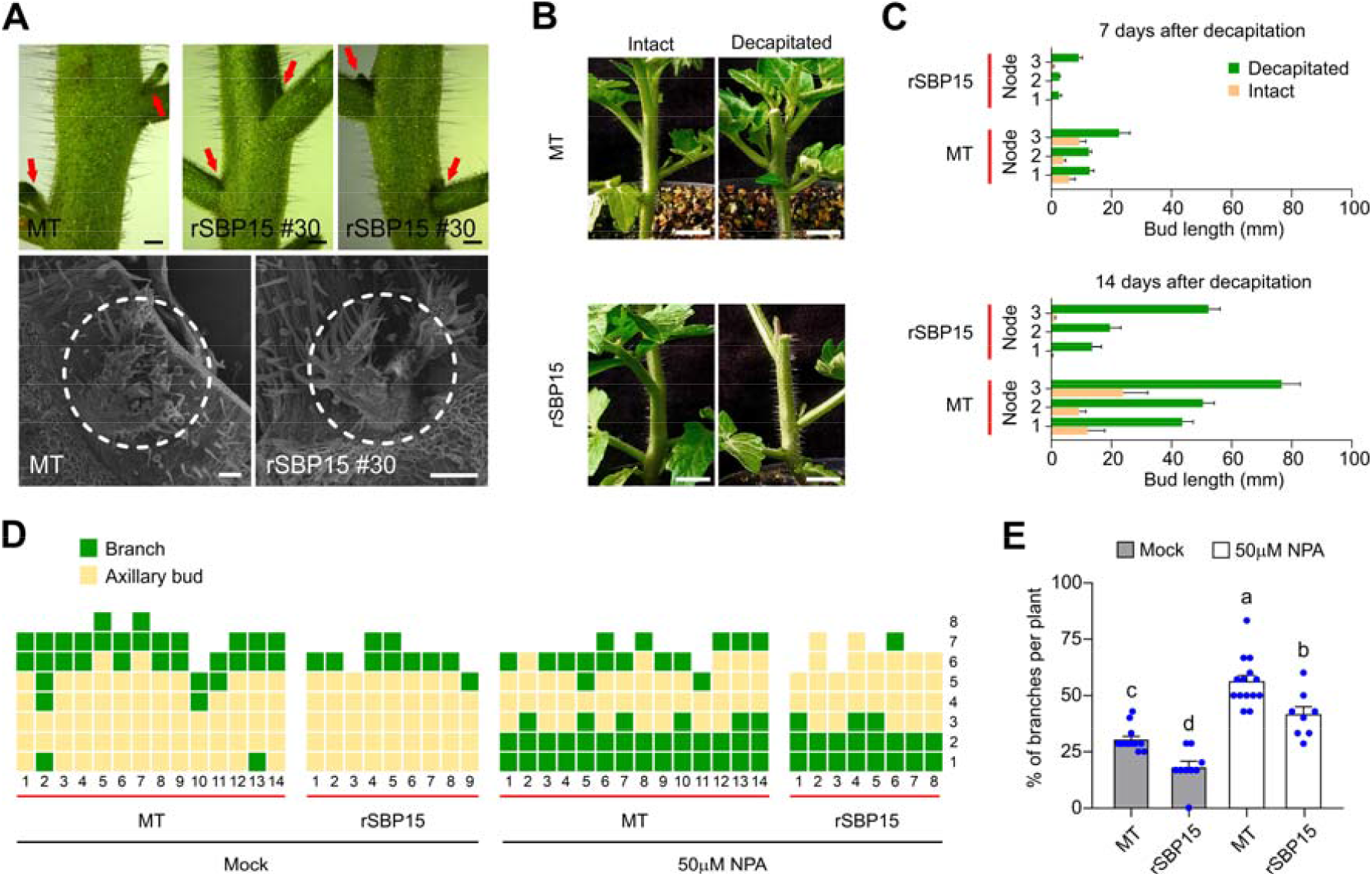
Loss of PAT triggers AB outgrowth in rSBP15 plants. (A) ABs from rSBP15 plants are smaller than MT (top, scale bars = 1 mm), which is confirmed by scanning electron microscope. Note the differences in the scale bar (bottom, scale bars = 200 µm). Red arrows and dashed white circles indicate ABs. (B) Representative pictures of intact and decapitated plants of MT and rSBP15 plants at 7 days after decapitation. Scale bars = 1 cm. (C) After decapitation, ABs were measured in intact (orange bars) and decapitated (dark green bars) plants after 7 (top) and 14 days (bottom). (D) SB patterns of MT and rSBP15 plants at 30-dpg treated with Mock and 50 µM NPA solutions. (E) Percentage of LBs per plant in MT and rSBP15 genotypes treated with Mock (gray bars) and 50 µM NPA (white bars) solutions. Values are mean ± SEM (n = 8-14). Distinct letters represent statistical differences at 5% level evaluated by Tukey’s HSD test. The blue dots represent individual data points. SB patterns were evaluated similarly as depicted in Figure 1.

### The miR156/*SlSBP15* hub affects auxin and ABA responses

ABs from rSBP15 plants are smaller and primarily dormant but are still able to outgrow after the loss of PAT. These observations suggest that the *SlSBP15* is involved in hormone responses associated to ABs dormancy/activation, however, the genes inside the ABs causing phytohormone responses remained unknown. To unravel the underlying phytohormones response-related molecular mechanisms within ABs, we have conducted NGS-based transcriptomics analyses (RNA-Seq) to compare the transcriptome in the ABs from 156-OE and rSBP15 plants at 25-dpg; at this time, we could harvest early/dormant ABs from all genotypes, thus the RNA-seq datasets were comparable. Samples were composed of at least 20 ABs from different plants. 439 and 2379 differentially expressed genes (DEGs) were identified in ABs from 156-OE and rSBP15 plants, respectively, compared to MT (Supplementary Tables S2; S3); in addition, we have identified 167 DEGs shared between 156-OE and rSBP15 ABs (Fig. 3A; Supplementary Table S4), being, sixty DEGs that showed opposite expression patterns in ABs from 156-OE and rSBP15 (Fig. 3B).

**Figure 3.**
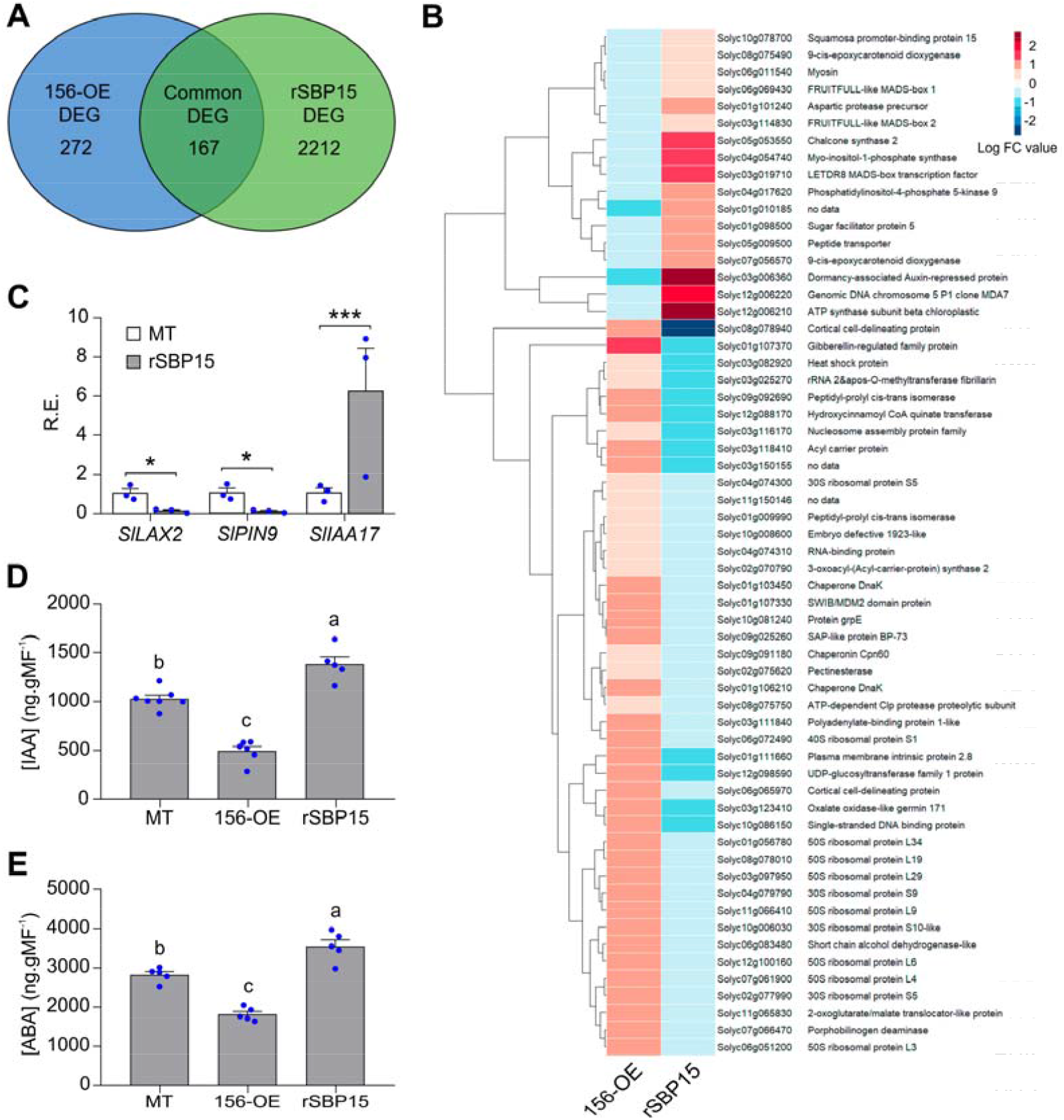
Auxin and ABA response are altered in early/dormant ABs from 156-OE and rSBP15 plants. (A) Venn diagram showing the number of common and unique DEGs between ABs from 156-OE and rSBP15 plants. DEG analyses were performed comparing either 156-OE or rSBP15 with MT expression data. (B) Expression profile of 60 DEGs that showed contrasting expression profiles between 156-OE and rSBP15 genotypes. The scale bar represents log2 fold-change values. (C) RE level by qRT-PCR of selected auxin-associated genes in samples of early/inactivate ABs from MT and rSBP15 plants. Values are mean ± SEM (n = 3). \**P*<0.05, \*\*\**P*<0.001, according to Student’s t-test (two-tailed). (D, E) IAA (D) and ABA (E) quantification in early/inactive ABs from MT, 156-OE and rSBP15 plants. Values are mean ± SEM. (n = 5-7). Distinct letters represent statistical differences at 5% level evaluated by Tukey’s HSD test. The blue dots represent individual data points.

Among these genes, several auxin-associated genes such as *SlLAX2* and *SlPIN9* involved in auxin distribution (Pattison and Catala, 2012), were down-regulated, and *SlIAA17*, a canonical Aux/IAA protein (Su et al., 2014), was strongly up-regulated in ABs from rSBP15 plants (Supplementary Table S3). To confirm this information, we have evaluated, by qRT-PCR analyzes the gene expression of *SlLAX2, SlPIN9* and *SlIAA17*, and have found similar results to those obtained by RNA-seq (Fig. 3C). These findings suggest that the ectopic expression of the resistant version of the *SlSBP15* to the miR156 cleavage led to defective auxin transport in ABs, which may result in IAA accumulation within ABs from rSBP15 plants. Thus, to further analyse, we have quantified, by gas chromatography-mass spectrometry (GC-MS) in selected ion monitoring (SIM), the IAA concentration in ABs from 156-OE and rSBP15 plants at 25-dpg, and we have found that IAA concentration was lower in ABs from 156-OE plants and higher in rSBP15 plants compared to MT (Fig. 3D), suggesting that the *SlSBP15-*mediated alteration in levels of auxin responses genes in tomato leads to dormancy of ABs from rSBP15 plants.

Our RNA-Seq analyses also revealed that several ABA-associated genes showed opposite expression patterns in ABs from 156-OE and rSBP15 plants; among them, we have identified two tomato *9-CIS-EPOXYCAROTENOID DIOXYGENASE-like* genes (*SlNCED* and *SlNCED-like*), key genes in ABA biosynthesis (Fig. 3B; Supplementary Table S4). Because of ABA acts as a negative regulator of AB activity (Yao and Finlayson, 2015) and *NCED* is a key enzyme in ABA biosynthesis, we have evaluated, by qRT-PCR analyzes, the gene expression of *SlNCED* in ABs from rSBP15 and 156-OE plants. *SlNCED* transcripts accumulated at higher levels in ABs from rSBP15 plants, but at lower levels in 156-OE plants (Supplementary Fig. S4A). This observation suggested that ABs from rSBP15 plants may accumulate ABA, which would correlate with their arrested development (Yao and Finlayson, 2015; González-Grandío *et al*., 2017). To confirm this information, we have also quantified, by GC-MS in SIM, the ABA concentration in ABs from 156-OE and rSBP15 plants at 25- dpg. Accordingly, low and high levels of ABA were observed in ABs from 156-OE and rSBP15 plants, respectively (Fig. 3E). Altogether, our results indicated that, although morphologically similar, early/dormant ABs from rSBP15 were still arrested, whereas those from MT and 156-OE plants had initiated their genetic reprogramming to outgrow.

### The miR156/*SlSBP15* hub affects dormancy-associated gene activity

So far, our data indicate that IAA and ABA concentrations are higher in ABs from rSBP15 plants compare to MT plants, suggesting that higher concentrations of IAA and ABA lead to ABs dormancy. The RNA-Seq analyses also revealed several dormancy-associated genes. Among them, *MADS-box* genes, known targets of SPLs/SBPs in several species, are associated with ABs arrest (Rosin *et al*., 2003; Wu *et al*., 2017). *SlTDR8, FRUITFULL-like MADS-box1* (*FUL1*) and *FUL2* were up-regulated in rSBP15, but down-regulated in 156-OE ABs (Supplementary Table S4; Fig. 3B; Supplementary Fig. S4B). Recently, *FUL1* and *FUL2* were shown to redundantly regulate inflorescence architecture in tomato. Null *ful1 ful2* tomato mutants showed high SB and delayed flowering (Jiang *et al*., 2022), similar to 156-OE plants (Fig. S1A-B). Thus, *SlTDR8, FUL1* and *FUL2* might be direct targets of SlSBP15 within ABs, although this prospect awaits further validation. In addition, we have found that early markers for ABs dormancy, such as a *DORMANCY-ASSOCIATED PROTEIN-LIKE1* gene (*SlDRM1*) (Stafstrom *et al*., 1998), were strongly down- and up-regulated in ABs from 156-OE and rSBP15 plants, respectively (Supplementary Table S4; Fig. 3B; Supplementary Fig. S4C). Other *DRM-like* genes (Solyc02g077880 and Solyc01g099840) were up-regulated only in ABs from rSBP15 (Supplementary Table S3). In contrast, transcript levels of the peptide-encoding *GIBBERELLIC ACID-STIMULATED ARABIDOPSIS* (*SlGASA*) were increased in 156-OE, but strongly reduced in ABs from rSBP15 plants (Supplementary Table S4; Fig. 3B; Supplementary Fig. S4 D). Interestingly, *Arabidopsis GASA6* was reported to function as an integrator of phytohormones gibberellin and ABA during seed germination (Zhong *et al*., 2015), whereas cotton *GhGASA10–1* gene is upregulated by IAA during fiber development (Chen et al, 2021), which indicates that GASA genes may integrate distinct hormonal pathways.

### The *SlSBP15* negatively regulates GOB at transcriptional level during SB

Our RNA-Seq analyses revealed that the miR164-targeted *GOB* was downregulated in ABs from both 156-OE and rSBP15 plants (Supplementary Tables S2, S3). *GOB* participates in tomato SB (Busch *et al*., 2011), and its activity is mediated by auxin in leaves (Ben-Gera *et al*., 2012). To further understand the relationship between GOB and the miR156/*SlSBP15* hub on ABs, we have evaluated, by qRT-PCR analyzes, the gene expression of GOB in ABs from 156-OE and rSBP15 plants, and confirmed the data obtained by RNA-seq. We have found that GOB expression was lower in ABs from both 156-OE and rSBP15 plants compared to MT (Fig. 4A), suggesting that the miR156/*SlSBP15* hub regulates *GOB* expression at the transcriptional level. To test this hypothesis, we have initially screened ∼3.8 kb promoter of *GOB* and found some possible *SPL*/*SBP* GTAC core motifs (Fig. 4B); then, we have generated the *pGOB::LUC* construct (∼3.8 kb *GOB* promoter fused to *LUCIFERASE*), and co-expressed it with *p35S::rSBP15*-harbouring vector, an empty vector, or *p35S::NLS-GFP*-harbouring vector in *N. benthamiana* leaves. Interestingly, Luciferase activity was significantly reduced in the presence of high levels of *rSBP15* (Fig. 4B). These observations suggested that SlSBP15 represses *GOB* expression within the ABs, which affects their competence to outgrowth. To investigate this possibility, we have crossed rSBP15 plants with the gain-of-function *GOB* mutant (*Gob-4d*) in MT background, which generates ectopic ABs and shows highly increased SB (Busch *et al*., 2011; Rossmann et al., 2015). As we expected, the rSBP15;*Gob-4d* plants display fewer number of active ABs and LBs compared to *Gob-4d* plants at 30 dpg (Fig. 4C, D), indicating that the miR156-resistant *SlSBP15* version is sufficient to arrest AB outgrowth in the highly branched *Gob-4d* plants.

**Figure 4.**
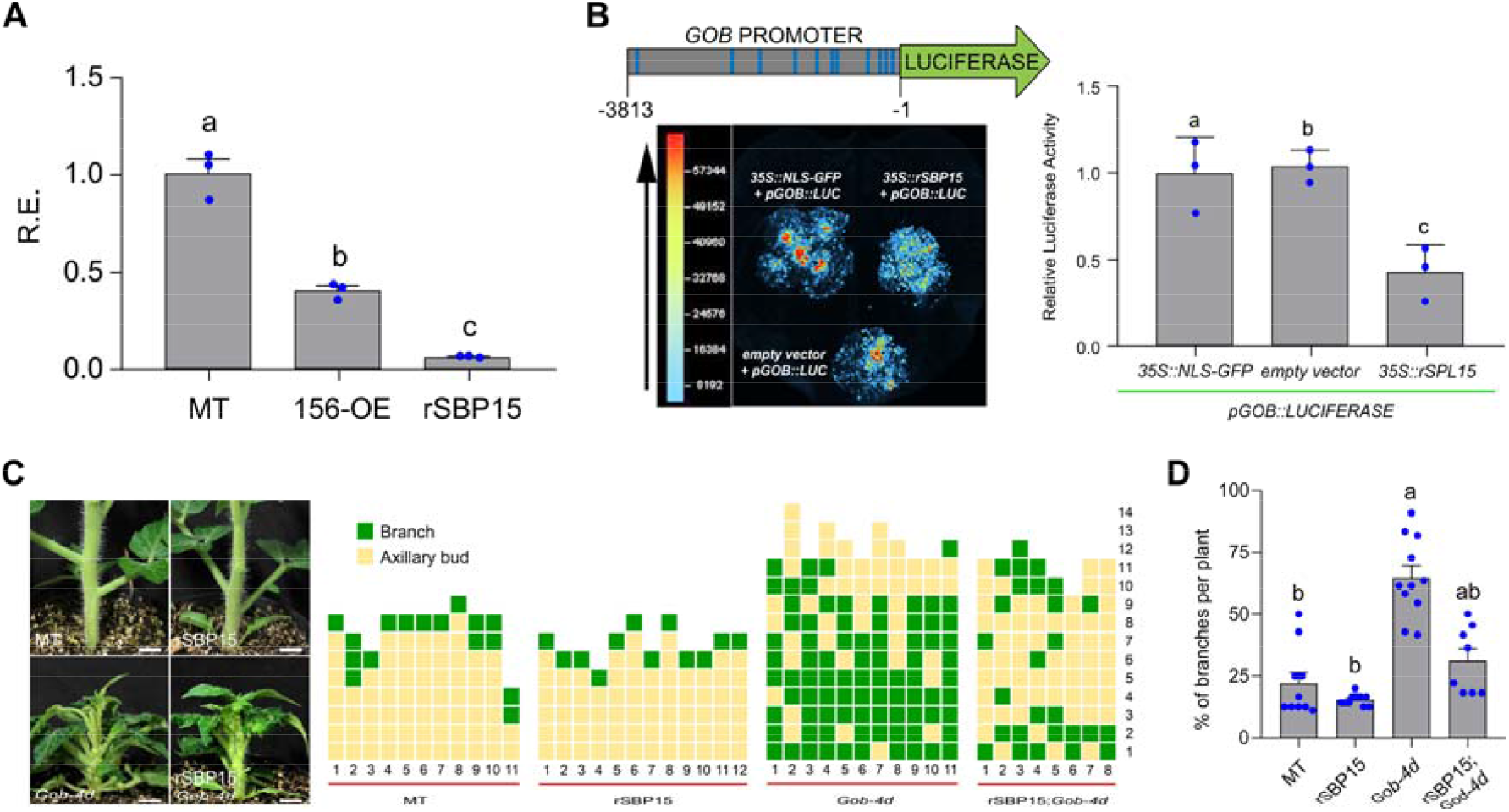
The *SlSBP15* regulates *GOBLET* (*GOB*) expression. **(A)** Relative expression level (R.E.) by qRT-PCR of *GOB* gene in early/inactive ABs of MT, 156-OE and rSBP15 plants. Values are mean ± SEM. (n = 3). Distinct letters represent statistical differences at 5% level evaluated by Tukey’s HSD test. **(B)** Regulation of the *GOB* promoter (*pGOB*) activity by rSBP15. Schematic representation of the 3813-bp *GOB* promoter region (gray box) fused to Luciferase (green box). Eleven cis-elements were identified as putative SBP binding sites (GTAC, blue lines in the gray box). *Nicotiana benthamiana* leaves were co-infiltrated with *Agrobacterium* containing *p35S::rSBP15, p35S::NLS-GFP* (control), empty vector (pK7WG2, control) and *pGOB::LUC* constructs. The luminescence images were captured using the NEWTON 7.0 CCD imaging system. Pseudocolors show the luminescence intensity (left). Quantification of relative luciferase activity (right) in leaves using NEWTON 7.0 imaging software. Values are mean ± SEM. (n = 3). Distinct letters represent statistical differences at 5% level evaluated by Tukey’s HSD test. **(C)** Representative pictures (left) and SB pattern (right) of MT, rSBP15, *God-4d*, and rSBP15;*Gob-4d* plants at 30-dpg. Scale bars = 1 cm. SB patterns were evaluated similarly as depicted in Figure 1. **(D)** Percentage of branches per plant in MT, rSBP15, *Gob-4d*, rSBP15;*Gob-4d* genotypes. Values are mean ± SEM (n = 8-12). Distinct letters represent statistical differences at 5% level evaluated by Dunn’s multiple comparisons test. The blue dots represent individual data points.

### SlSBP15 interacts with SlBRC1b to regulate downstream targets

Homologues of *SlSBP15* in wheat, rice and *Arabidopsis* (Supplementary Fig. 1D) inhibit SB, at least in part, by activating the expression of *TB1-*like genes (Lu *et al*., 2013; Liu *et al*., 2017; Xie *et al*., 2020). Tomato *TB1-*like genes *BRC1a* (Solyc03g119770) and *BRC1b* (Solyc02g089830) are expressed in arrested ABs and both are down-regulated upon ABs activation (Martín-Trillo et al, 2011). However, we found neither *SlBRC1a* nor *SlBRC1b* in our transcriptomic data (Supplementary Tables S2-S3). We then hypothesized that the *SlSBP15* interacts with the *SlBRC1* at the protein level. To test this possibility, we conducted yeast two-hybrid (Y2H) assays between SlSBP15 and SlBRC1a/b proteins. SlSBP15 interacted with SlBRC1b but not with SlBRC1a, as shown by yeast growth in selective media (Fig. 5A). To further confirm this interaction, we performed Acceptor Photo-Bleaching Fluorescence Resonance Energy Transfer (APB–FRET) assay of fluorescent protein fusions of SlBRC1b co-expressed with a miR156-resistant version of SlSBP15 (rSBP15) in *Nicotiana* leaves. The E_FRET_ (percentage of GFP expression change) values for SlBRC1b:GFP/rSBP15:mCherry were not as high as for SlBRC1b:GFP/SlBRC1b:mCherry, but they were significantly higher than SlBRC1b:GFP, a negative control (Fig. 5B). These findings demonstrated that SlSBP15 specifically interacts with SlBRC1b and probably modulates its transcriptional activity towards common targets.

**Figure 5.**
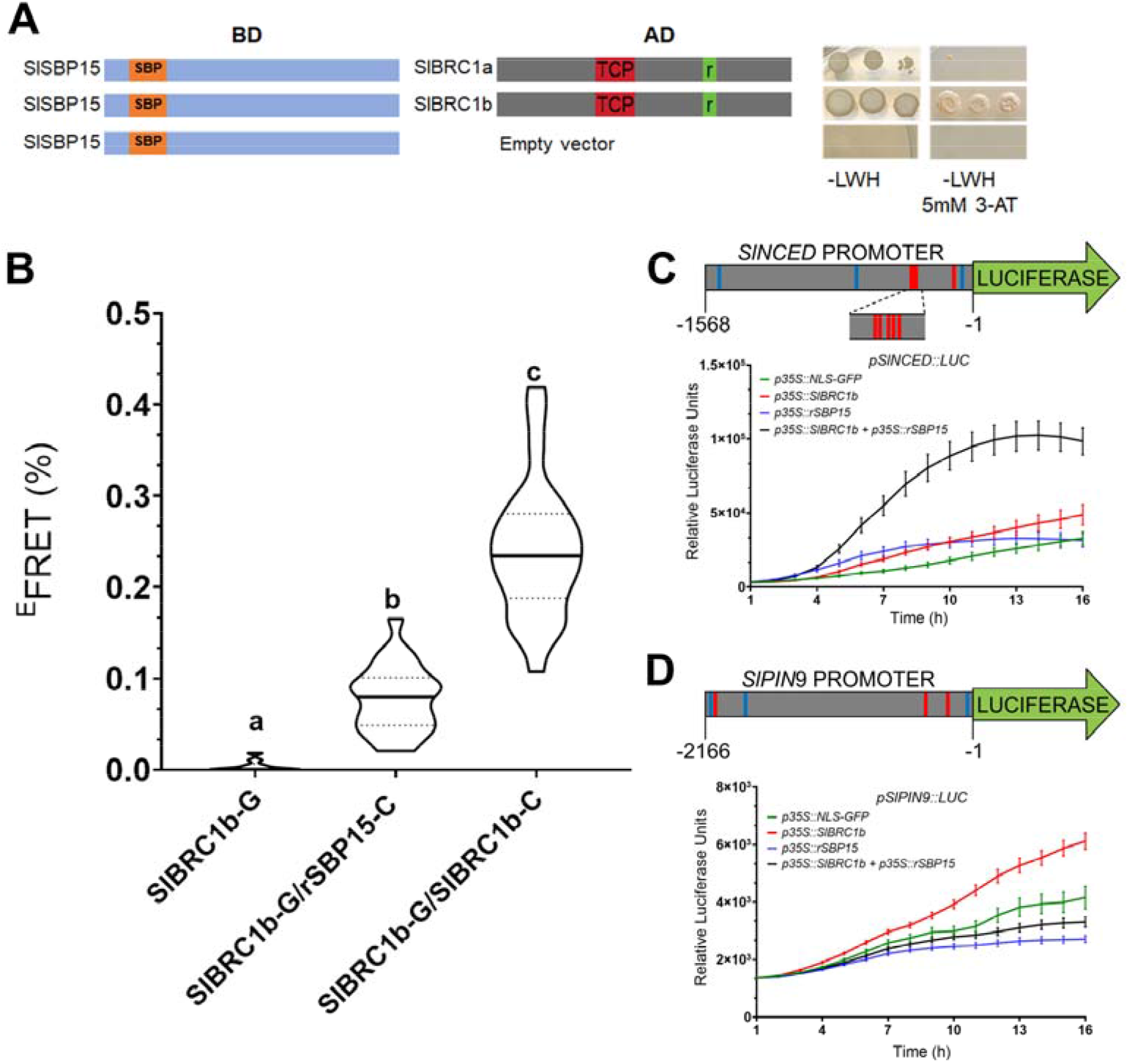
SlSBP15 acts in concern and independently of SlBRC1b to control tomato AB out growth. **(A)** SlSBP15 and SlBRC1b interact in a yeast-two-hybrid assay. SlSBP15 ORF was cloned into pGBKT7-BD vector, while SlBRC1b ORF (including the TCP and R domain; Cubas et al., 1999) was cloned in pGADT7-AD. Vectors were co-transformed in the yeast strain AH109. Transformants were selected in media lacking leucine and tryptophan (-LH) and assayed for interaction in media lacking leucine, tryptophan and histidine (-LWH) in the presence or absence of 5mM of 3-amino-1, 2, 4-triazole (3-AT). Each row shows ten-fold serial dilutions of the indicated strain. **(B)** FRET acceptor photobleaching measurements between SlBRC1b and rSBP15. FRET efficiency (^E^FRET) is calculated as relative increase in GFP (G) fluorescence intensity after photobleaching of the mCherry (C) acceptor. Intramolecular FRET (positive control with both fluorophores in the same protein, SlBRC1b-G/C) and negative control (donor only, SlBRC1b-G) are included. Results show the mean ± SEM. n = 10-25 cells from at least four different leaves. Distinct letters represent statistical differences at 0.1% level evaluated by Tukey’s HSD test. **(C, D)**. Luciferase (LUC) assays showing the activity of *pSlNCED1::LUC* (**C**) and *pSlPIN9::LUC* (**D**) when co-infiltrated with *p35S::NLS-GFP* (control), *p35S::SlBRC1b*, and *p35S::rSBP15* constructs. Schematic representation of *SlNCED* (1568 bp) and *SlPIN9* (2166 bp) promoter region (gray box) fused to Luciferase (green box). Cis-elements were identified as putative SBP (blue lines) and TCP (red lines) binding sites.

In *Arabidopsis, BRC1* enhances *NCED3* expression within ABs (González-Grandío *et al*., 2017), whereas cucumber *BRC1* (*CsBRC1*) directly represses the expression of *CsPIN3* (Shen *et al*., 2019). Based on these reports, we have investigated in detail *SlNCED* and *SlPIN9*, two possible direct targets of *SlSBP15* identified in our transcriptomics, and both confirmed by qRT-PCR (Supplementary Tables S2-S4; Fig. 3C; Supplementary Fig. S4A). Screening for BRC1-like and SBP-like consensus binding sites (González-Grandío *et al*., 2017; Van Es *et al*., 2020; Wang and Wang, 2015) revealed partially conserved core motifs in the *SlNCED* and *SlPIN9* promoter regions (∼1.6 kb and ∼2.2 Kb, respectively) for both proteins (Supplementary Fig. S5). To test the *SlNCED* and *SlPIN9* response to SlSBP15 and SlBRC1b, we co-expressed a *LUC* reporter fused to either promoter (*pSlNCED::LUC* and *pSlPIN9::LUC*) with either the presence of *p35S:rSBP15* or *p35S:SlBRC1b*, or both constructs. We monitored LUCIFERASE activity for 16 hours in *Nicotiana* leaves. For the *pSlNCED* promoter, LUCIFERASE activity increased in several time-points in the presence of either rSBP15 or SlBRC1b (Fig. 5C). Strikingly, SlBRC1b and rSBP15 cooperatively enhanced *pSlNCED::LUC* activity at much higher levels relative to the other treatments. On the other hand, *pSlPIN9::LUC* activity was activated by SlBRC1b, but repressed by rSBP15 (Fig. 5D). The combination of SlBRC1b and rSBP15 led to a weak repression of *SlPIN9* promoter activity, suggesting that SlSBP15 antagonizes or acts independently of SlBRC1b in the context of *SlPIN9* regulation. These results indicated that tomato BRC1b is sufficient to modulate SlNCED and SlPIN9 expression not only in ABs, where SlBRC1b is mostly expressed, but also in leaves where SlBRC1b usually is low expressed (Martín-Trillo et al., 2011). Most importantly, our data indicated that SlSBP15 and SlBRC1b can modulate the expression of common targets cooperatively and antagonistically.

## Discussion

In cultivated tomato, SB is an undesirable trait as it competes for photoassimilates with developing flowers and fruits. Thus, manipulating genes associated with increasing apical dominance could help to generate improved tomato architecture with reduced axillary shoots. Here, we have shown that, not only the establishment of tomato vegetative architecture is dependent on the miR156 because of overexpression of this highly conserved miRNA triggers SB as it happens in Arabidopsis, rice, maize, and wheat (Gou et al., 2017; Liu *et al*., 2017; Luo *et al*., 2012; Chuck *et al*., 2007; Schwab *et al*., 2005); but also, we have demostrated that the miR156-targeted *SlSBP15* interconnects hormones such as auxin and ABA with major SB-associated genes such us *GOB* and *SlBRC1b* forming a regulation circuit of SB; thus, the *SlSBP15*, or even its downstream targets, may be important candidates for manipulating this specific aspect in tomato breeding programs.

Our phylogenetic analysis of *SPL*s/*SBP*s genes has demostrated that, among the SBP tomato genes, only the *SlSBP15* grouped into the cluster containing known SB-associated SPL genes from other especies, indicating that the *SlSBP15* regulates the SB in tomato. When ectopically expressed, the miR156-resistant *SlSBP15* (rSBP15 plants) has a strong AB-arresting activity in distinct cultivated tomatoes (Fig. 1C, D; Supplementary Fig. S3); however, the WT-like branching pattern of *sbp15*^*CRISPR*^ mutants (Fig. 1E, F) sugests that the *SlSBP15* plays redundant roles with other miR156-targeted *SlSBPs* within the SB context. This points out a possible scenario in which *SlSBP15* is functionally redundant with *SlSBP13* in the context of SB (Cui *et al*., 2020). Thus, future generation of *sbp13*^*CRISPR*^ *sbp15*^*CRISPR*^ double mutants will help to address whether stronger branching phenotypes can be obtained by knocking out both *SlSBP* genes.

Auxin is mainly produced in the SAM and transported basipetally mostly through the PAT stream. To outgrow, ABs need to export their auxin (Li and Bangerth, 1999); interestingly, basal ABs in rSBP15 plants responded to the loss of PAT stream, but not the apical ones (Supplementary Fig. S2D, E). The altered auxin transport observed in rSBP15 plants is, at least in part, caused by the mis-regulation of auxin efflux protein-encoding genes, such as *PIN9*, and *LAX2* (Fig. 3C; Supplementary Table S3); Pattison and Catala, 2012) suggesting that the *SlSBP15* regulates *PIN* transporters, as occurs in rice, where IPA1 directly binds to the promoter of *OsPIN1b* and activates its transcription (Lu et al., 2013); however, it remains to be demonstrated whether miR156-targeted SlSBPs directly bind to *PIN* and *LAX* promoters to regulate their activities.

Auxin modulates the activity of transcription factors associated with the control of SB, such as *GOB*, a fundamental fundamental factor for establishing tomato AMs (Ben-Gera *et al*., 2012; Busch *et al*., 2011), and it is regulated by the *SlSBP15* module (Supplementary Table S3; Fig. 4A, B; Silva *et al*., 2014). Our data indicate that low levels of *GOB* transcripts may be crucial to maintain ABs from rSBP15 at the early/dormant stage, as the overexpression of *SlSBP15* resistant-version in the highly branched *Gob-4d* drastically reduces the number of active ABs (Fig. 4 C, D). It is possible that auxin-mediated *GOB* repression (Ben-Gera *et al*., 2012) is controlled by the *SlSBP15*-dependent regulation within ABs. Given that LS transcript accumulation is dependent on GOB activity (Rossmann et al., 2015), and no LS transcripts were observed in our transcriptome analysis (Supplementary Tables S1; S2), the SlSBP15 likely regulates LS and GOB at distinct AM and AB developmental stages..

*IPA1* and its orthologs in wheat and *Arabidopsis* have been reported to act upstream of *TB1*/*BRC1* genes by directly activating their expression (Lu *et al*., 2013; Li *et al*., 2017; Xie *et al*., 2020). Here, we have shown that the SlSBP15 role in ABs arrest is not mediated by *SlBRC1b* activation, but rather SlSBP15 and SlBRC1b interact at the protein level. Interactions between SPLs/SBPs and members of the TCP family have already been described. In *Arabidopsis*, for example, SPL9 interacts with TCP4 (Rubio-Somoza *et al*., 2014), and IPA1 can interact with PCF1 and PCF2 in rice (Lu *et al*., 2013). Importantly, SlSBP15 and SlBRC1b seem to inhibit tomato SB by cooperatively modulating downstream targets. One such example is the activation of *SlNCED* by both SlSBP15 and SlBRC1b (Fig. 5C), which lead to high levels of ABA content in dormant ABs from rSBP15 plants. Within ABs from *Arabidopsis, BRC1*, together with HD-ZIP-encoding genes such as *HOMEOBOX PROTEIN 40* (*HB40*), enhances *SlNCED3* expression, leading to ABA accumulation and suppression of ABs development (González-Grandío *et al*., 2017). We observed nearly a two-fold induction of the tomato homolog of *HB40* in early/dormant ABs from rSBP15 (Supplementary Table S3). Thus, we can envision a molecular circuity comprising *SlSBP15-SlBRC1b-HB40-SlNCED* that modulates ABA levels within tomato ABs.

Similar genetic factors control flowering time and SB in tomato, including the miR156/*SlSBP* hub (Weng *et al*., 2016; Silva *et al*., 2019). Thus, manipulating the transcription of genes associated with both developmental processes has great potential applications in breeding programs. Our findings that high levels of *SlSBP15* negatively regulates SB suggest that manipulation of this gene might be an important tool in molecular breeding for obtaining new tomato cultivars. With the emergence of novel gene editing techniques, fine-tune controlling the activity of the miR156/*SlSBP* hub should be further explored as a target to achieve the theoretical “IDEAL TOMATO ARCHITECTURE” by simultaneously modulating flowering time and SB.

## Acknowledgments

We thank the members of Dr Nogueira’s laboratory for helpful discussions. We are in debt to Prof. Lazaro Peres (ESALQ/University of Sao Paulo, Brazil) for providing seeds of the *Gob-4d* mutant and wild tomato relatives. We thank the members of Dr. Pilar Cubas group (CNB/CSIC, Spain) for providing technical help and insightful discussions.

## Author contributions

C.H.B-R., D.A.P.B, M.H.V. and F.T.S.N. were responsible for the conception, planning and organization of the experimental time line. C.H.B-R., D.A.P.B, M.H.V., E.M.S., A.M.L., L.F.F., G.F.F.S., J.P.O.C. carried out experiments. F.T.S.N., L.F., and P.C. directly supervised the development of the experimental plan. R.M.C. and C.M.S.S. helped to analyze the RNA-seq data. C.H.B-R., D.A.P.B, M.H.V. and F.T.S.N. discussed the resulting data. The manuscript was written by C.H.B-R., M.H.V. and F.T.S.N.

---

**Supplementary Figure S1.**
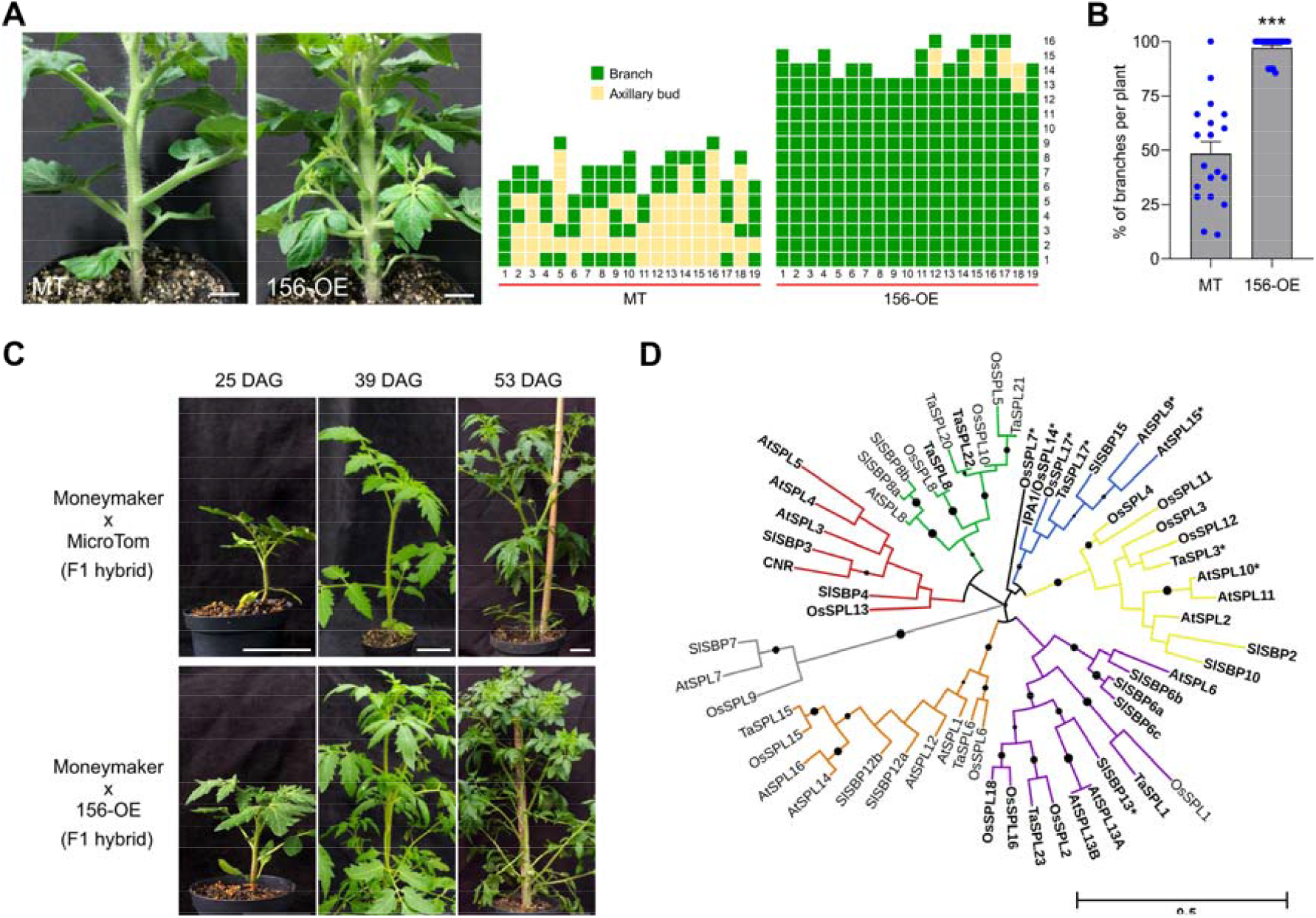
miR156-targeted *SlSBPs* modulate tomato SB. (**A**) Representative pictures (left) and SB patterns (right) of MT and 156-OE plants at 30-dpg. Each column represents a single plant, and each square indicates the status of each leaf axil starting from the first node above the cotyledons. AB are shown in orange squares and LB in dark green squares. (**B**) Percentage of LB per plant in 156-OE and MT at 30-dpg. The percentage was determined based on the number of leaf axils that present LB. Values are mean ± SEM (n = 19). \*\*\**P*<0.001, according to Mann Whitney test. The blue dots represent individual data points. **C**. The LB phenotype of 156-OE plants is not background dependent. Representative phenotype of F1 offspring resulting from the crossing between 156-OE and MT with tomato cv. Moneymaker plants at 25, 39 and 53-dpg. Scale bars = 5 cm. **D**. Phylogenetic analysis of SQUAMOSA PROMOTER-BINDING PROTEIN LIKE (SPLs/SBPs) in *Arabidopsis* (AtSPLs), rice (OsSPLs), wheat (TaSPLs), and tomato (SlSBPs). The SBP domains were aligned using ClustalW and the tree was constructed using the maximum likelihood algorithm with 1000 bootstraps. Nodes supported by bootstrap values higher than 50% are marked with a dot, proportional to the bootstrap value. miR156-targeted SPLs/SBPs are in bold. Different clades are indicated with different colors. Asterisks indicate SPLs previously associated with SB. Red arrow indicates *SlSBP15*.

**Supplementary Figure S2.**
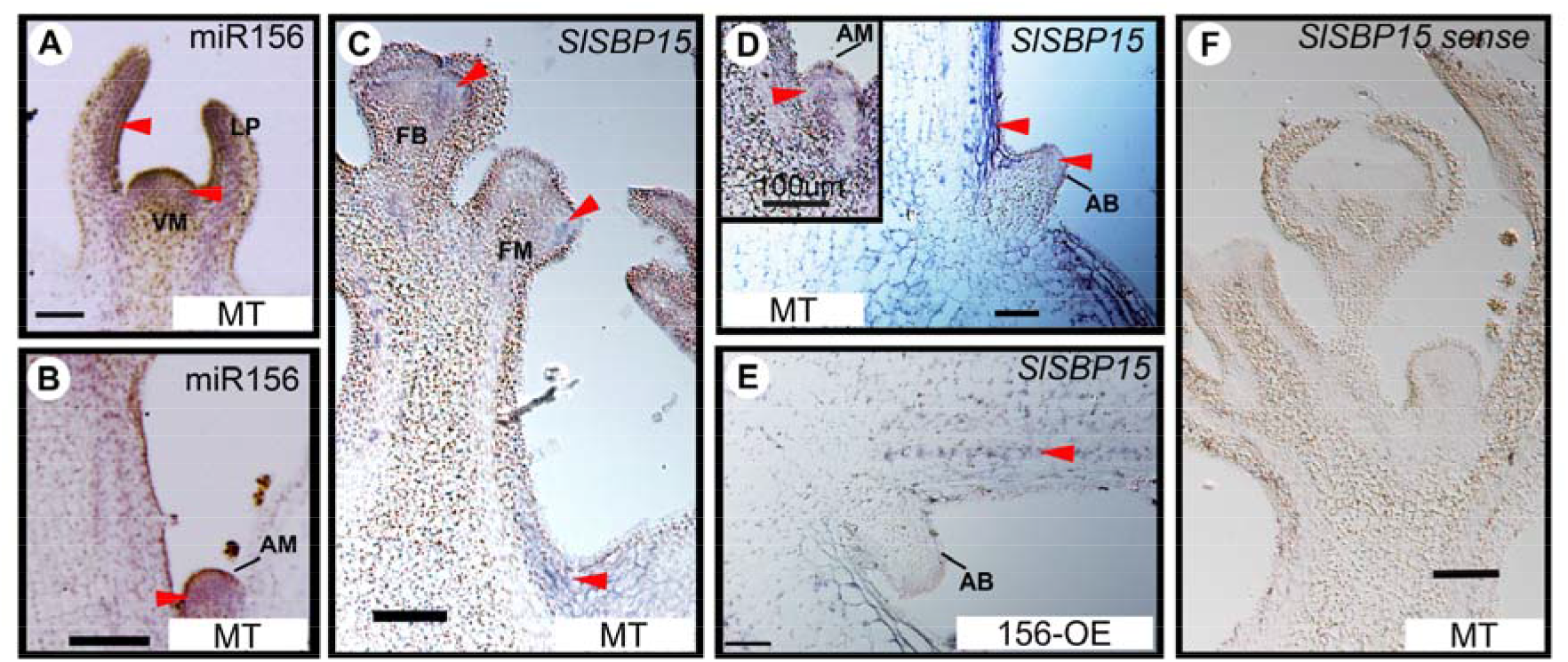
Expression patterns of miR156 and *SlSBP15* in tomato apexes. **(A-B)** *In situ* hybridization showing the miR156 expression in leaf primordia (LP), SAM (VM, vegetative meristem) and AM in tomato seedlings at 7- and 10-dpg. **(C-E)**. A digoxigenin-labeled probe detecting *SlSBP15* transcripts was hybridized with longitudinal sections of MT and 156-OE apexes. **(C)** *SlSBP15* transcripts were detected in the floral bud (FB), floral meristem (FM), and leaf axil. **(D)** *In situ* hybridization showing *SlSBP15* expression in AB, AM (inset), and subepidermal tissues close to AB. **(E)** In 156-OE seedlings, *SlPBP15* expression was restricted to the procambium. **(F)** *In situ* hybridization of *SlPBP15* sense probe was used as a negative control. Purple staining shows probe localization (arrowheads). Scale bars = 100 µm.

**Supplementary Figure S3.**
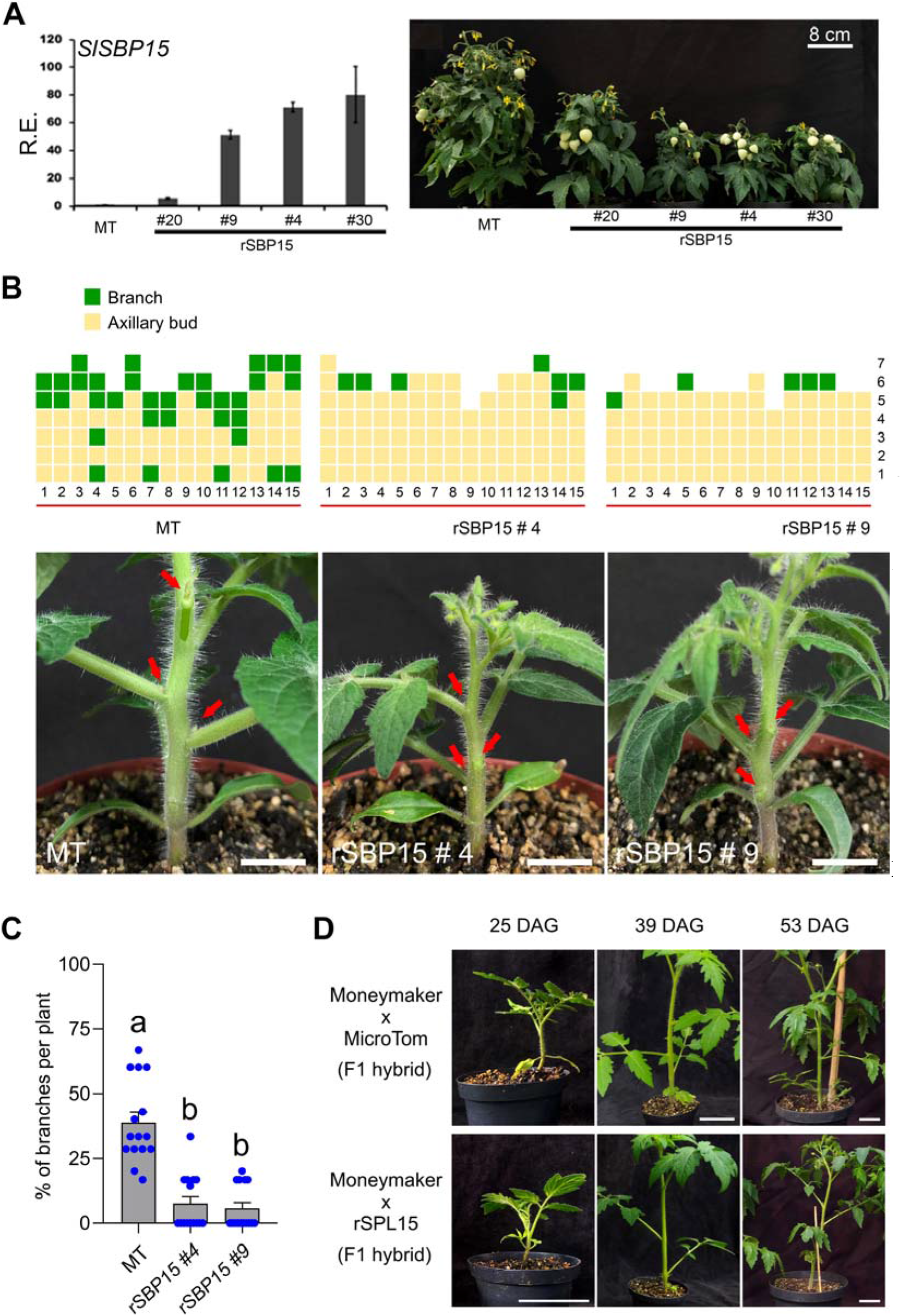
Molecular and phenotypic characterization of rSBP15 transgenic line. **(A)** Relative expression (R.E.) by QRT–PCR of *SlSBP15* transcript levels in leaves and representative pictures of MT, and rSBP15 lines #20, #4, #9 and #30. Values are mean ± SEM. (n = 3). **(B)** SB pattern (top) and representative pictures (bottom) of MT, and rSBP15 lines #4 and #9 at 30-dpg. Each column represents a single plant and each square indicates the status of each leaf axil starting from the first node above the cotyledons. ABs (orange squares) or LBs (dark green squares). Red arrows indicate ABs. Scale bars = 1 cm. **(C)** Percentage of LBs per plant in MT and rSBP15 (lines #4 and # 9) genotypes, at 30-dpg. The percentage was determined based on the number of leaf axils that present LBs. Values are mean ± SEM (n = 15). Distinct letters represent statistical differences at 5% level evaluated by Tukey’s HSD test. The blue dots represent individual data points. **(D)** Representative phenotype of F1 offspring resulting from the crossing between MT and rSBP15 with tomato cv. MM plants at 25, 39 and 53-dpg. Red arrowheads indicate leaf axils. Scale bars = 5 cm.

**Supplementary Figure S4.**
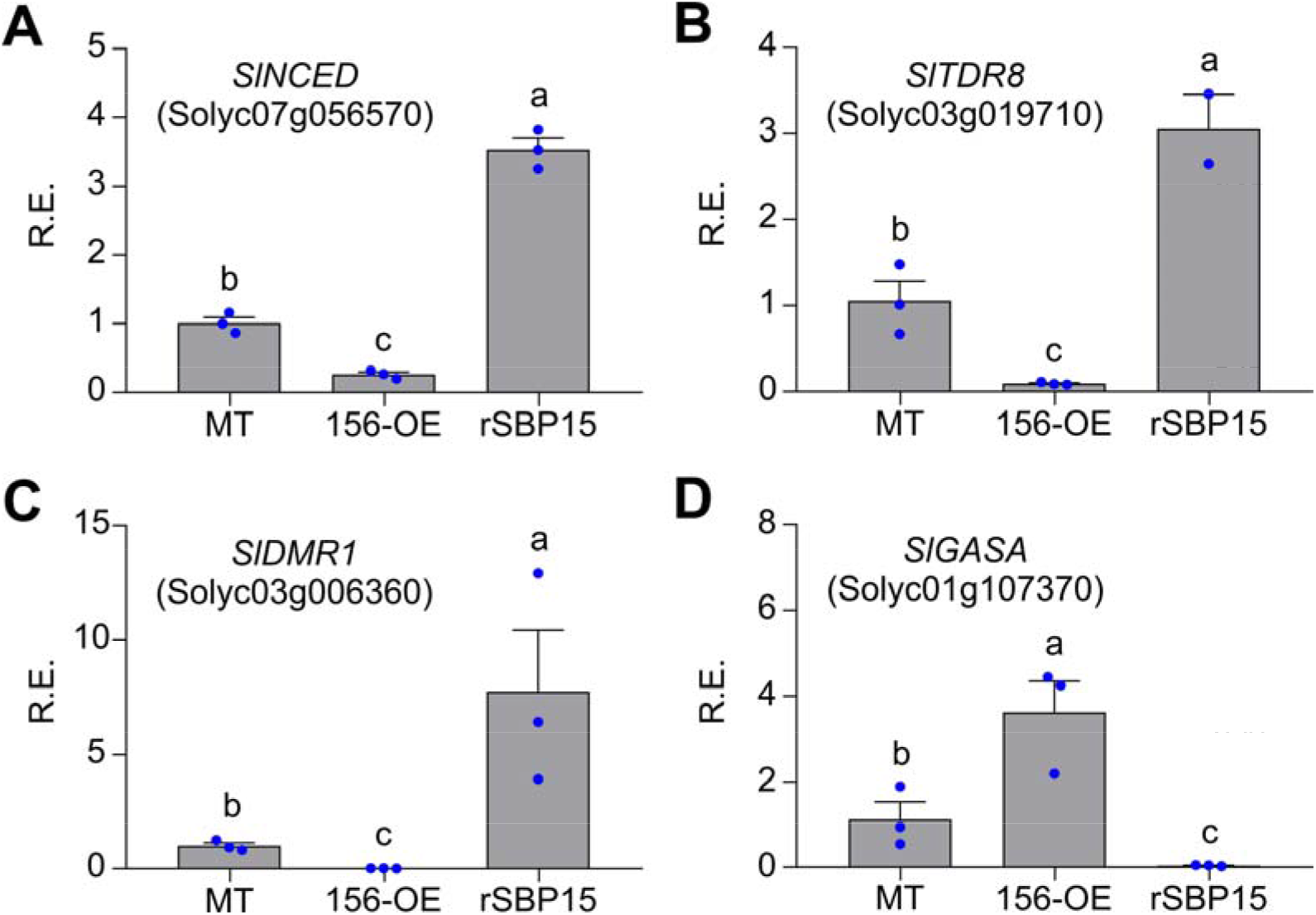
The miR156/SISBP15 hub modulates early markers for ABs dormancy. **(A-D)** RE level of *SlNCDE1* (**A**), *SlTDR8* (**B**), *SlDRM1* (**C**), and *SlGASA* (**D**) by qRT-PCR in independent samples of MT, 156-OE, and rSBP15 early/inactive ABs. *SlNCDE1, 9-cis-epoxycarotenoid dioxygenase*; *SlTDR8, MADS-box transcription factor*; *SlDRM1, DORMANCY-ASSOCIATED PROTEIN 1*; and *SlGASA, Gibberellin-regulated family protein*;. Values are mean ± SEM. (n = 3). Distinct letters represent statistical differences at 5% level evaluated by Tukey’s HSD test. The blue dots represent individual data points.

**Supplementary Figure S5.**
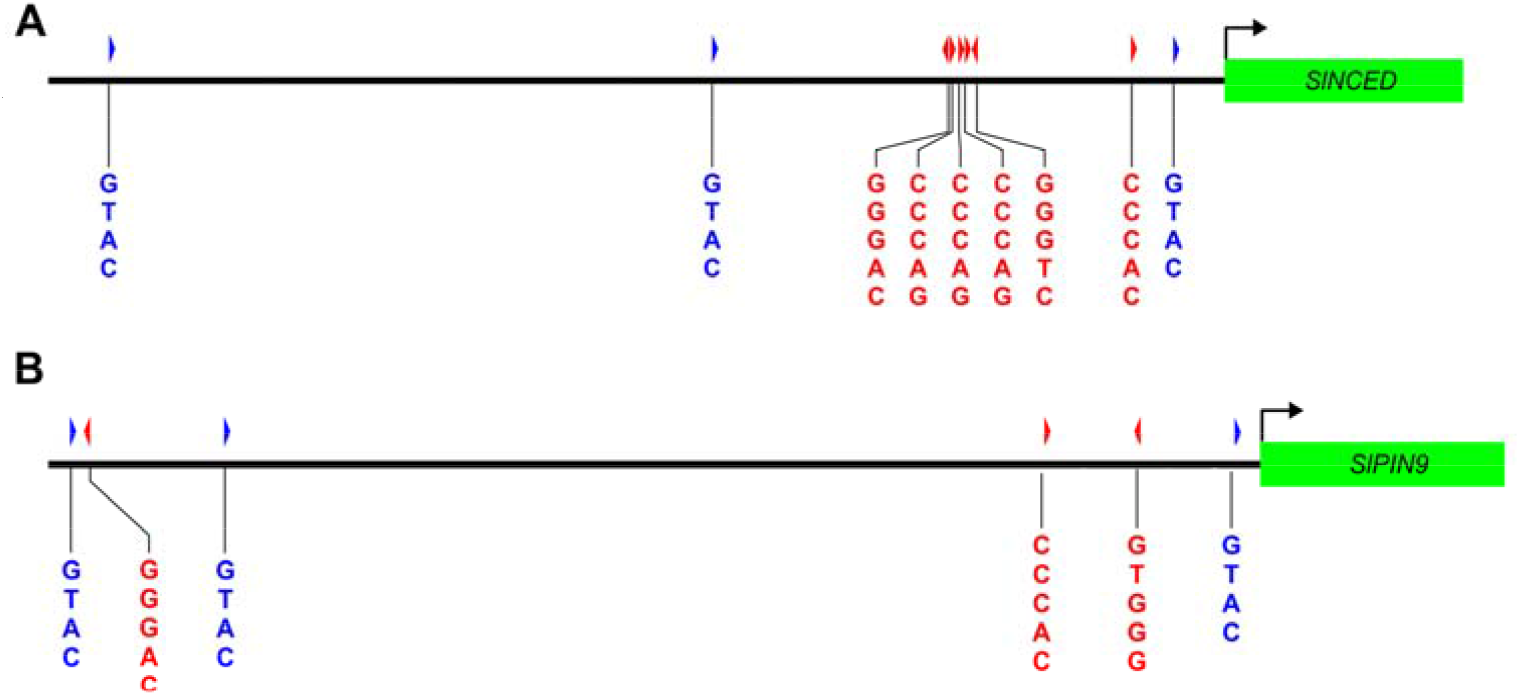
Representation of *SlNCED* and *SlPIN9* promoters. (A) Schematic diagram of the 1568-bp *SlNCED* promoter region. Three and six cis-elements were identified as putative SBP (blue sequence) and TCP (red sequence) binding sites, respectively. (B) Schematic diagram of the 2166-bp *SlPIN9* promoter region. Three putative binding sites were identified for SBP (blue sequences and arrowheads) and three for TCP (red sequences and arrowheads). GTAC binding motif is recognized by SPLs/SBPs, while GnCCC and CCCAC are recognized by TCPs. Putative core sequence motifs are presented in the 5’-to 3’-orientation. Black line and green box indicate promoter and coding regions, respectively.

